# Designing mRNA coding sequence via multimodal reverse translation language modeling with Pro2RNA

**DOI:** 10.64898/2026.03.18.712790

**Authors:** Bian Bian, Yiming Zhang, Jichen Zhang, Kiyoshi Asai, Yutaka Saito

**Author notes:** Correspondence authors: Bian Bian, Yutaka Saito. Equally contributed (Bian Bian, Yiming Zhang, Jichen Zhang).

## Abstract

mRNA coding sequence design is a critical component in the development of mRNA vaccines, nucleic acid therapeutics, and heterologous gene expression systems. While large language models have recently been successfully applied to protein design and RNA modeling, designing optimal mRNA coding sequences for a given protein, particularly in a species-specific manner, remains a major challenge. Here, we present Pro2RNA, a multimodal reverse-translation language model that generates mRNA coding sequences from their corresponding protein sequences while explicitly conditioning on host organism taxonomy information. Pro2RNA integrates multiple pretrained language models across different modalities, including ESM2 for protein representation, SciBERT for taxonomy understanding, and a generative RNA language model for mRNA codon-level sequence generation. By training on mRNA-protein pairs from eukaryote and bacteria datasets respectively, Pro2RNA learns species-dependent genetic codes and codon usage patterns, enabling the generation of host-adapted and natural-like mRNA coding sequences. Across multiple benchmark evaluations, Pro2RNA matches or surpasses existing optimization methods, demonstrating its potential as a powerful and flexible framework for species-aware mRNA coding sequence design.

## Introduction

The design of mRNA coding sequences is a critical step in the development of mRNA vaccines, nucleic acid therapeutics, synthetic biology, and heterologous gene expression systems (Jin et al. 2025; Paremskaia et al. 2024; Pfeifer et al. 2023; Zhang et al. 2025b). Owing to the degeneracy of the genetic code, multiple synonymous codons can encode the same amino acid. However, codon usage is far from uniform: different organisms exhibit distinct codon preferences shaped by evolutionary pressure, tRNA abundance, and translational regulation (Kober and Pogson 2013; Hershberg and Petrov 2008; Plotkin and Kudla 2011). Consequently, effective mRNA coding sequence design must account for host-specific features to achieve optimal translation efficiency and protein expression.

Traditional mRNA coding sequence optimization strategies primarily rely on host-specific codon usage tables, replacing rare codons with so-called “optimal” synonymous codons (Schmidt et al. 2023; Tian et al. 2017; Sharp and Li 1987). While this approach can increase overall translation rates, it neglects the global sequence context of genes (Mazumdar et al. 2017). Increasing evidence suggests that rare codons are not merely translational bottlenecks; instead, they can play functional roles in regulating ribosome pausing and co-translational protein folding (Moss et al. 2024; Bian et al. 2023; Komar 2009). Disrupting these patterns through naive codon replacement may lead to misfolded proteins, reduced functional yield, or impaired biological activity (Yu et al. 2015). These observations indicate that effective mRNA design requires learning deeper, organism-specific sequence rules that go beyond simple codon frequency statistics.

The emergence of large language models (LLMs) has transformed our ability to model complex biological sequences (Sarumi and Heider 2024; Lin et al. 2025; Liu et al. 2025). In natural language processing, pretrained language models such as SciBERT and BioBERT leverage large-scale scientific and biomedical corpora to capture semantic, syntactic, and contextual information, enabling robust representation learning for downstream tasks (Beltagy et al. 2019; Lee et al. 2020). Analogously, biological language models trained on large-scale sequence data have demonstrated the ability to capture hierarchical and contextual features of biomolecules (Brixi et al. 2025; Nguyen et al. 2024). In the protein domain, transformer-based models such as the ESM series, built on BERT-like architectures, learn rich representations of protein sequences that encode structural, functional, and evolutionary information (Hayes et al. 2025; Lin et al. 2023). In parallel, generative architectures such as GPT-style and Hyena-based models have further expanded sequence modeling capabilities by enabling long-range dependency modeling and autoregressive generation (Ferruz et al. 2022; Zhang et al. 2025c). In the RNA domain, similar advances have been achieved. Encoder-based models such as RNA-FM apply BERT-style architectures to non-coding RNA sequences, capturing conserved structural and functional patterns (Chen et al. 2022). Generative models such as GenerRNA further demonstrate the potential of language models for non-coding RNA sequence design (Zhao et al. 2024). More recently, models specifically targeting mRNA coding sequences have emerged, including encoder-based approaches such as CodonBERT and NUWA, as well as generative frameworks such as mRNA-GPT (Li et al. 2024; Bian et al. 2025; Zhong et al. 2025). These models move beyond handcrafted features and enable data-driven learning of codon usage, regulatory elements, and global sequence context, providing deeper insights into mRNA biology.

The design of mRNA coding sequences can be naturally formulated as a reverse translation problem, in which an mRNA sequence is generated from a given protein sequence under specific host constraints. In this study, we introduce Pro2RNA, a multimodal protein-to-mRNA reverse translation language model that integrates pretrained biological language models to enable host-aware mRNA sequence generation and optimization. Pro2RNA combines a protein language model (ESM2) and natural language model (SciBERT) as encoder module, together with a generative mRNA language model (mRNA-GPT&GenerRNA) as the decoder, explicitly incorporating host taxonomy information to guide mRNA sequence generation. By leveraging the rich representations learned by large-scale pretrained biological language models, Pro2RNA provides a unified and flexible framework for mRNA coding sequence design, with broad application in biomanufacturing, synthetic biology, and mRNA vaccine development.

## Results

### Pro2RNA integrated pretrained sciBERT, ESM2 and generative RNA language model for mRNA coding sequence design

Pro2RNA is designed to generate optimized mRNA coding sequences conditioned on protein sequences and host taxonomy information. To evaluate the impact of different architectural components, we conducted a systematic ablation study by combining pretrained SciBERT and ESM2 as encoder and RNA language models with different decoder modules (Figure1).

Specifically, we compared four model variants: ESM2+SciBERT+MLP, ESM2 + SciBERT + GenerRNA, ESM2 + SciBERT + mRNA-GPT, and ESM2 + mRNA-GPT. For all configurations, we adopted a consistent training strategy using LoRA fine-tuning and trained each model on data from three bacterial species (Table 1A). Model performance was evaluated on the same held-out test dataset using the naturalness score, which measures statistical consistency with species-specific codon usage patterns.

**Table 1.** Architecture variants of the Pro2RNA framework and their performance across different model combinations. **(A)** Representative bacterial species used to evaluate different architectural combinations of the Pro2RNA framework. RefSeq IDs and the corresponding scientific names are listed. **(B)** Performance comparison of different Pro2RNA model variants with respect to Naturalness score, together with the number of trainable parameters.

| RefSeq ID | Scientific Name |
| --- | --- |
| GCF_964059955.1 | *Sodalis-like secondary symbiont of Drepanosiphum platanoidis* |
| GCF_964062745.1 | *Levilactobacillus cerevisiae* |
| GCF_964063155.1 | *Pediococcus parvulus* |

**B**
| Model structure | Naturalness score | Trainable parameters |
| --- | --- | --- |
| Pro2RNA (ESM2+SciBERT+GenerRNA) | 0.1264 | 24.5M |
| Pro2RNA (ESM2+SciBERT+MLP) | 0.0122 | 14.6M |
| Pro2RNA (ESM2+SciBERT+mRNA-GPT) | 0.5506 | 24.5M |
| Pro2RNA (ESM2+mRNA-GPT) | 0.5301 | 21.9M |

As shown in Table 1B, the ESM2 + SciBERT + mRNA-GPT configuration achieved the highest naturalness score (0.5506), substantially outperforming architectures based on MLP decoders or GenerRNA. This result is expected, as mRNA-GPT is pretrained on coding mRNA sequences, whereas GenerRNA is trained on non-coding RNAs and therefore lacks codon-level generation capacity, leading to markedly lower naturalness scores.

Moreover, incorporating SciBERT improved model performance compared with the variant without SciBERT (ESM2 + mRNA-GPT; overall naturalness score = 0.5301), the full model achieved a higher overall naturalness score (0.5506) (Table 1B) and consistently outperformed the ablated model across all three bacterial species (Table 2). This improvement suggests that biological text semantics captured by SciBERT provide complementary, taxonomy-aware contextual information beyond protein sequence embeddings, enabling the model to generate mRNA coding sequences that are more species-consistent and biologically plausible.

**Table 2.**
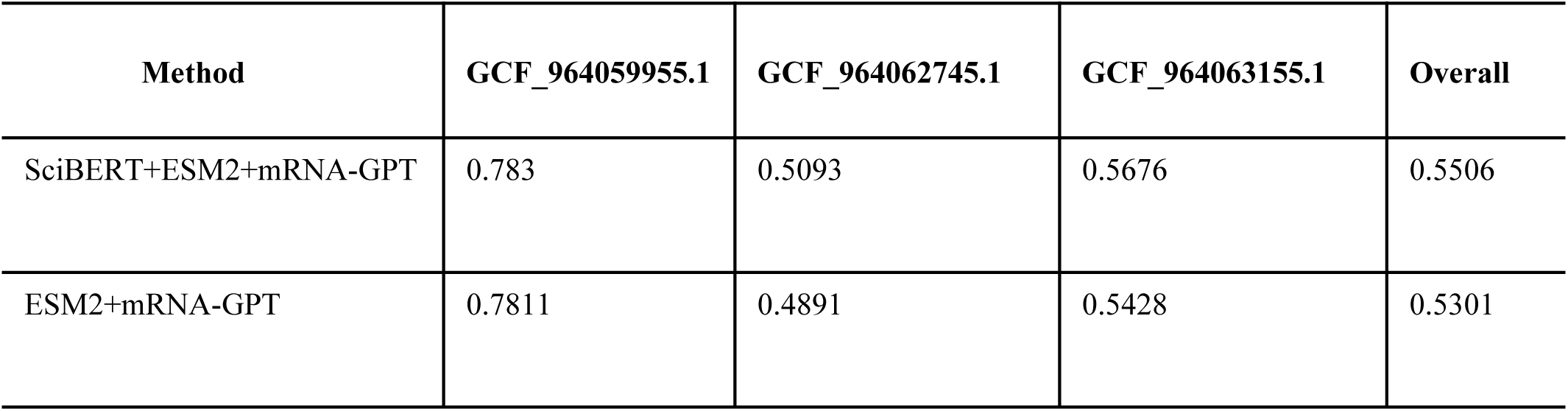
Ablation study of Pro2RNA assessing the contribution of taxonomy encoding across bacterial species. Performance comparison of Pro2RNA variants with and without the SciBERT based taxonomy encoder on three bacterial species (RefSeq IDs: GCF_964059955.1, GCF_964062745.1, and GCF_964063155.1). Results are reported using the Naturalness score,

| Method | GCF_964059955.1 | GCF_964062745.1 | GCF_964063155.1 | Overall |
| --- | --- | --- | --- | --- |
| SciBERT+ESM2+mRNA-GPT | 0.783 | 0.5093 | 0.5676 | 0.5506 |
| ESM2+mRNA-GPT | 0.7811 | 0.4891 | 0.5428 | 0.5301 |

Based on these results, we designate the ESM2 + SciBERT + mRNA-GPT architecture as Pro2RNA and use it for all subsequent training and analyses.

### Pro2RNA-eukaryote learned the genetic code and coding patterns in eukaryote

To evaluate whether Pro2RNA can learn eukaryotic codon usage patterns, we first selected 13 representative eukaryotic organisms (Table S1) and trained a model referred to as Pro2RNA-eukaryote. We initially assessed the average naturalness score across these species and observed consistently high values, indicating that the model effectively captures eukaryote-specific codon usage characteristics (Table 3).

**Table 3.** Cross-domain performance of Pro2RNA models trained on different organism groups. Comparison of Pro2RNA variants trained on different organismal domains, including eukaryotes, human-specific data, and bacteria. Performance is evaluated using the Naturalness score across multiple test settings: the average of 13 eukaryotic species, human, Candida albicans, and the average of 261 bacterial species.

| Method | 13 eukaryotes<br>(Average) | human | *C.albicans* | 261 bacteria<br>(Average) |
| --- | --- | --- | --- | --- |
| Pro2RNA-eukaryote | 0.5391 | 0.5438 | 0.5748 | 0.3718 |
| Pro2RNA-human | 0.5026 | 0.5318 | 0.4923 | 0.4539 |
| Pro2RNA-bacteria | 0.4199 | 0.4424 | 0.5318 | 0.5866 |

To further examine whether border-species training improves model performance, we compared Pro2RNA-eukaryote with a model trained exclusively on human data (Pro2RNA-human). The results show that Pro2RNA-eukaryote achieves a higher naturalness score than Pro2RNA-human in human test dataset, demonstrating that training on multiple related species enhances codon usage modeling beyond single-species training (Table3). Furthermore, we evaluated both models on completely unseen eukaryotic species. Pro2RNA-eukaryote consistently generated mRNA coding sequences with high naturalness scores (0.5748), whereas the single-species model (Pro2RNA-human) showed reduced performance (0.4923) in *C.albicans* (Table 3). These results indicate that border-species training substantially improves generalization to unseen eukaryotic species, enabling Pro2RNA to produce biologically plausible and species-consistent mRNA sequences.

In addition, we compare Pro2RNA-eukaryote with CodonTransformer, and three widely used vendor algorithm: Twist Bioscience, Integrated DNA technology (IDT), and Genewiz which focused on four eukaryotic model organism: *Saccharomyces cerevisiae*, *Arabidopsis thaliana*, *Mus musculus*, and *Homo sapiens*. To ensure a fair comparison, we evaluate all methods on the same protein sets in the CodonTransformer paper (Fallahpour et al. 2025). To evaluate the generated codon sequences with the natural codon sequence, we used the following evaluation metrics: codon similarity score which is the codon-level identity fraction between the generated and ground-truth codon sequences, codon similarity index (CSI) is defined as the geometric mean of species-specific codon weights along a sequence. In addition, we choose the dynamic time warping (DTW) distance which quantifies the similarity between two sequences by allowing flexible, non-linear alignment to account for local shifts and variations. Pro2RNA achieve highest codon similarity score and highest CSI value across all the organisms compare to other method. In addition, mRNA coding sequences generated by Pro2RNA-eukaryote attains low DTW distance to the ground truth sequence across most organism (Figure 2). Together, these results support that Pro2RNA-eukaryote outperformed existed method in eukaryotic hosts which captures the natural multi-scale structure of codon suage pattern, from local codon composition to global bias.

**Figure 1.**
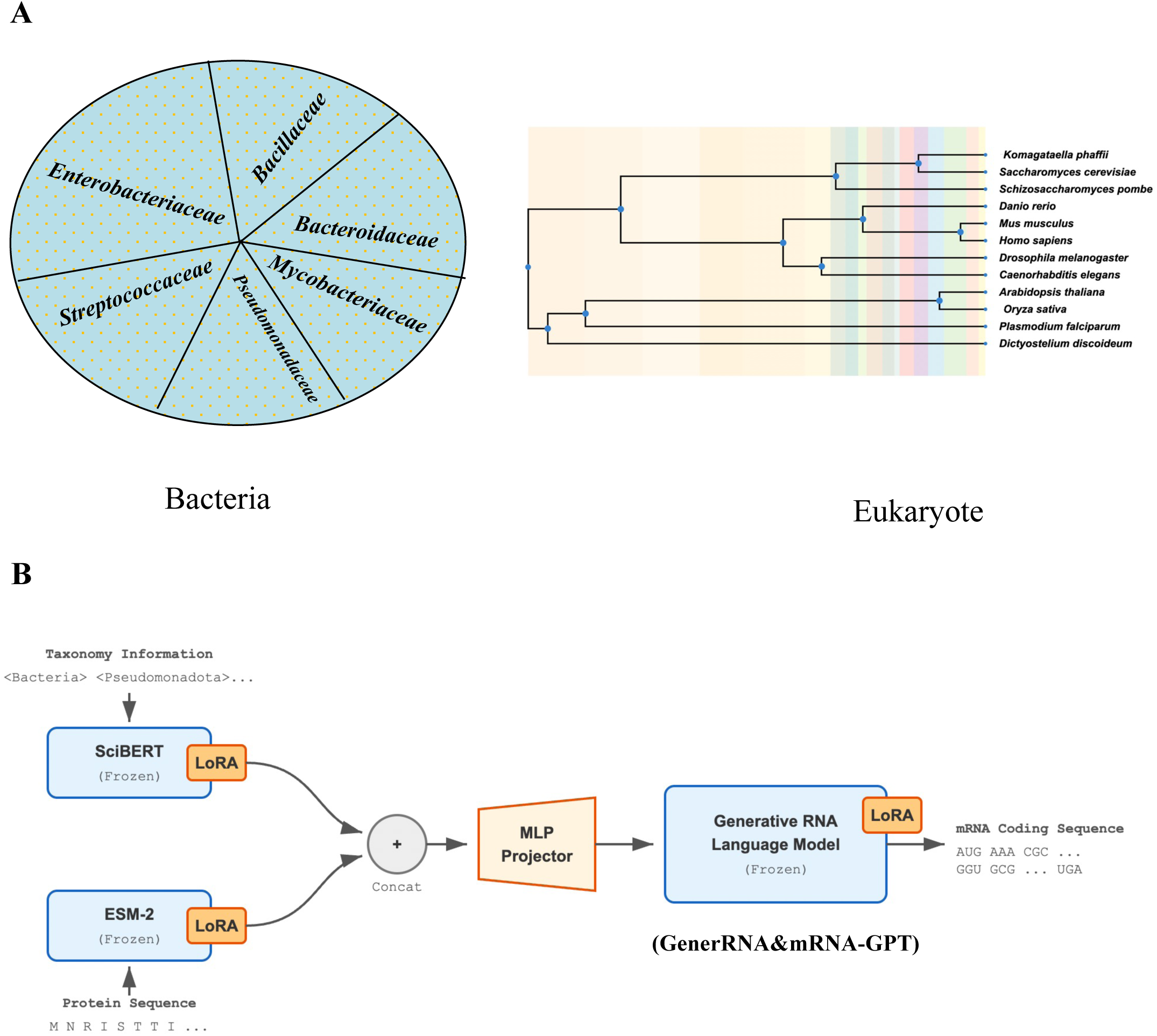
Overview of the Pro2RNA framework for mRNA coding sequence design. (A) Landscape of the training dataset, covering diverse bacterial and eukaryotic taxa. The bacterial dataset spans multiple representative families, while the eukaryotic dataset includes species distributed across major evolutionary lineages. (B) Schematic of the Pro2RNA architecture. Taxonomic information (e.g., domain, phylum, order, and family) is encoded using a pretrained language model (SciBERT), while the input protein sequence is encoded by a pretrained protein language model (ESM-2). Both encoders are kept frozen, with lightweight Low-Rank Adaptation (LoRA) modules introduced to enable efficient task-specific adaptation. The taxonomy and protein representations are concatenated and projected through an MLP module, and subsequently fed into a generative RNA language model (GenerRNA / mRNA-GPT), where LoRA is also applied, to generate species-specific mRNA coding sequences in a codon-level manner.Frozen parameters are shown in blue, while trainable components, including LoRA and the projector, are highlighted in orange.

**Figure 2.**
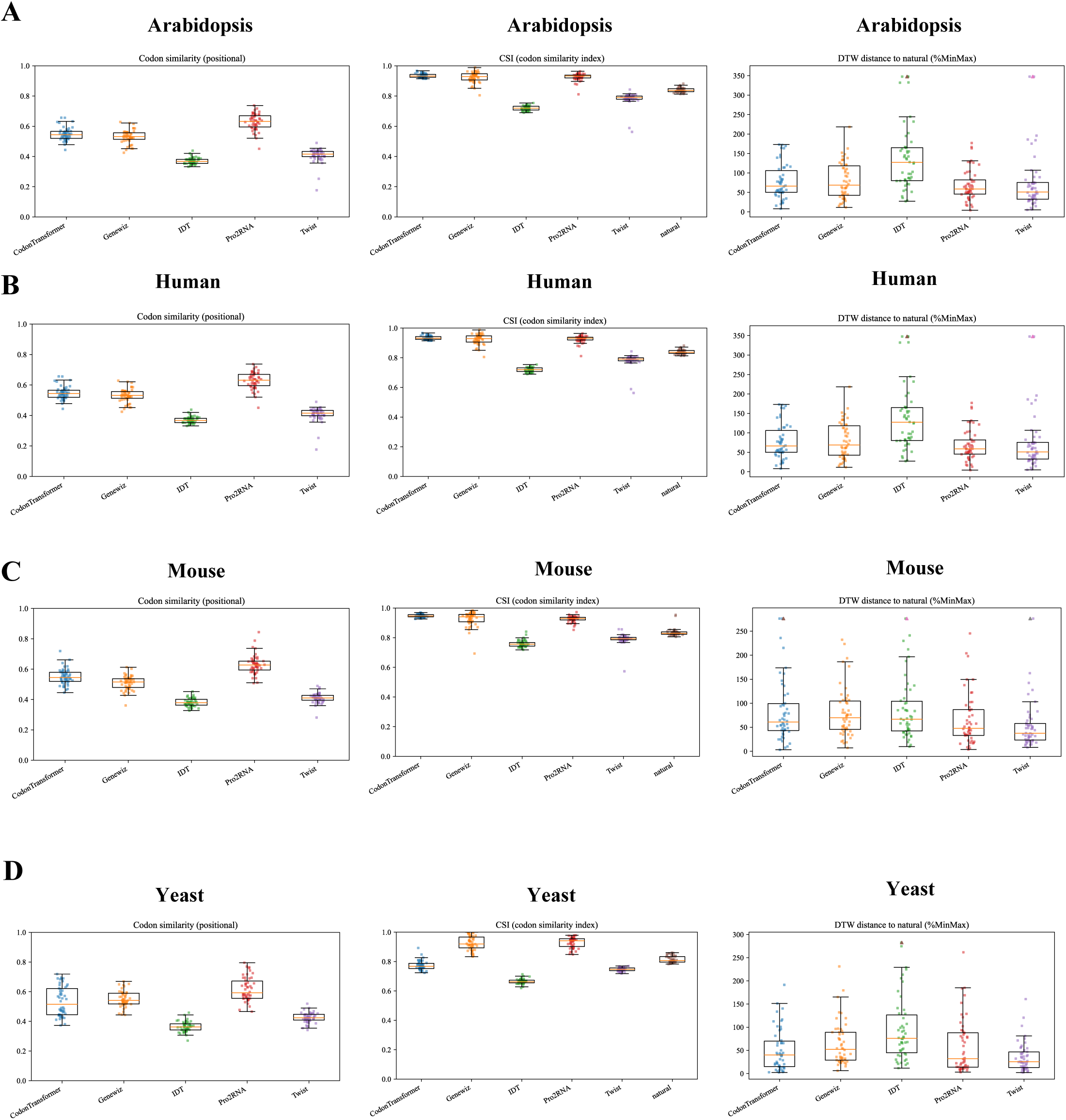
Pro2RNA-eukaryote captures organism-specific coding patterns and generates natural-like coding sequences across eukaryotes. Comparative evaluation of five codon optimization strategies on four representative eukaryotic model organisms: (A) Arabidopsis thaliana, (B) Human, (C) Mouse, and (D) Yeast. The naturalness of generated coding sequences relative to native sequences was assessed using three metrics: codon similarity, Codon Similarity Index (CSI), and Dynamic Time Warping (DTW) distance of normalized %MinMax profiles. Higher codon similarity and CSI values indicate closer agreement with native codon usage patterns, whereas lower DTW distances reflect greater similarity to natural translational dynamics. Box plots summarize the distribution of metrics across test genes, with individual dots representing single sequences.

### Pro2RNA-bacteria learned the genetic code and coding patterns in bacteria

Bacteria serve as the primary hosts for heterologous protein expression, and many classical bacterial species play essential roles in biomanufacturing and microbial cell factories (Gustafsson et al. 2004; Lee et al. 2009). Consequently, optimizing eukaryotic mRNA coding sequences for efficient protein expression in bacterial systems is a critical step (Kudla et al. 2009; Rosano and Ceccarelli 2014). Directly expressing native mRNA codon sequences from higher eukaryotes in bacteria often fails; even when expression is achieved, protein yield is frequently low, or the resulting proteins are misfolded (Komar 2009; Zhou et al. 2013).

To learn bacterial mRNA coding patterns and enable the rational design of mRNA coding sequences for bacterial expression, we constructed a large-scale bacterial training dataset spanning six representative families: *Bacillaceae*, *Bacteroidaceae*, *Enterobacteriaceae*, *Mycobacteriaceae*, *Pseudomonadaceae*, and *Streptococcaceae* (Table S2). In total, the dataset comprises 261 bacterial species and nearly one million paired mRNA-protein sequences. Model training was conducted on eight NVIDIA H200 GPUs for 15 epochs spent 23 hours. The loss curve of Pro2RNA-bacteria decreased monotonically and remained stable throughout training, indicating healthy optimization dynamics and strong generalization capability (Figure S1). On the held-out test dataset, the model achieved a naturalness score of 0.5866 (Table 3).

To benchmark Pro2RNA-bacteria against existing methods, we used the same protein dataset in CodonTransformer paper (Fallahpour et al. 2025) and evaluated performance using the same metrics applied in the eukaryotic setting. Pro2RNA-bacteria consistently outperformed other methods in terms of codon similarity score and codon similarity index (CSI), while achieving comparable performance in dynamic time warping (DTW) in *E.coli* (Figure 3A). These results demonstrate that Pro2RNA-bacteria successfully captures species-specific bacterial mRNA codon encoding rules, enabling the design of mRNA coding sequences optimized for bacterial heterologous expression.

**Figure 3.**
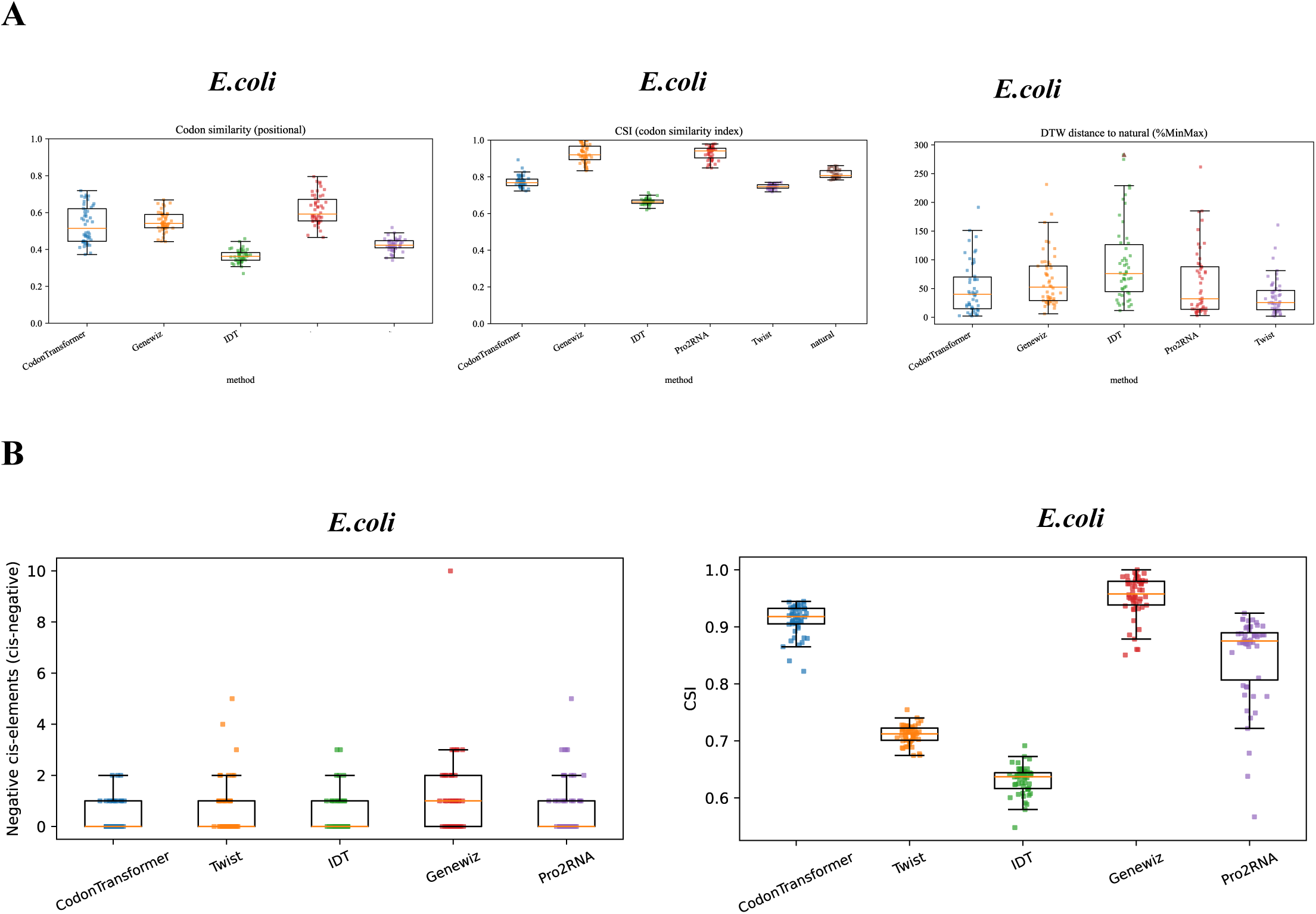
Pro2RNA-bacteria captures species-specific coding patterns and improves heterologous expression in *E. coli*. (A) Comparative evaluation of five codon optimization strategies in *E. coli*, assessing how well generated coding sequences recapitulate native E. coli codon usage patterns. Three metrics were used: codon similarity, Codon Similarity Index (CSI), and Dynamic Time Warping (DTW) distance of normalized %MinMax profiles. Higher codon similarity and CSI values indicate better agreement with endogenous coding patterns, while lower DTW distances reflect closer alignment with native translational dynamics. (B) Performance of different codon optimization methods on heterologous (exogenous) protein expression in *E. coli*. Left, distribution of negative cis-regulatory elements (CIS-negative) within generated coding sequences. Right, CSI scores of optimized sequences.

### Pro2RNA generate coding sequence with reduced negative *cis*-regulatory elements and intermediate CSI for heterologous expression

To systematically benchmark mRNA coding sequence design performance under heterologous expression settings, we selected 52 representative proteins curated from previous studies and designed corresponding mRNA coding sequences for expression in five host organisms spanning both prokaryotes and eukaryotes, including *Escherichia coli*, *Saccharomyces cerevisiae*, *Arabidopsis thaliana*, *Mus musculus*, and *Homo sapiens*. For each designed mRNA coding sequence, we quantified codon preference alignment using the codon similarity index (CSI) and assessed the abundance of predicted negative *cis*-regulatory elements within the coding sequence.

Across all evaluated hosts, Pro2RNA consistently generated mRNA sequences with reduced numbers of negative *cis*-regulatory elements, while maintaining intermediate CSI values relative to other methods (Figure 3B and Figure 4). In contrast, several existing approaches, particularly commercial pipeline Genewiz, tended to maximize CSI aggressively, often yielding sequences with extremely high codon preference alignment but simultaneously introducing a higher burden of negative *cis*-elements (Figure 3B and Figure 4). Although higher CSI values generally indicate stronger alignment with host-preferred codon usage, over-optimization toward highly preferred codons can be detrimental, as it may increase local mRNA secondary structure, disrupt beneficial ribosome stalling, or interfere with co-translational protein folding. Notably, Pro2RNA avoids indiscriminate CSI maximization and instead produces sequences whose CSI distributions are intermediate, suggesting that the Pro2RNA captures species-specific codon usage patterns rather than enforcing uniform optimization rules. This behavior is consistent across all tested organisms. In mammalian and plant hosts, Pro2RNA exhibits CSI values intermediate between highly optimized commercial designs, accompanied by consistently lower predicted negative *cis*-regulatory elements. These results indicate that Pro2RNA balances codon preference alignment with regulatory sequence integrity, producing mRNA designs that are both host-aware and structurally restrained.

**Figure 4.**
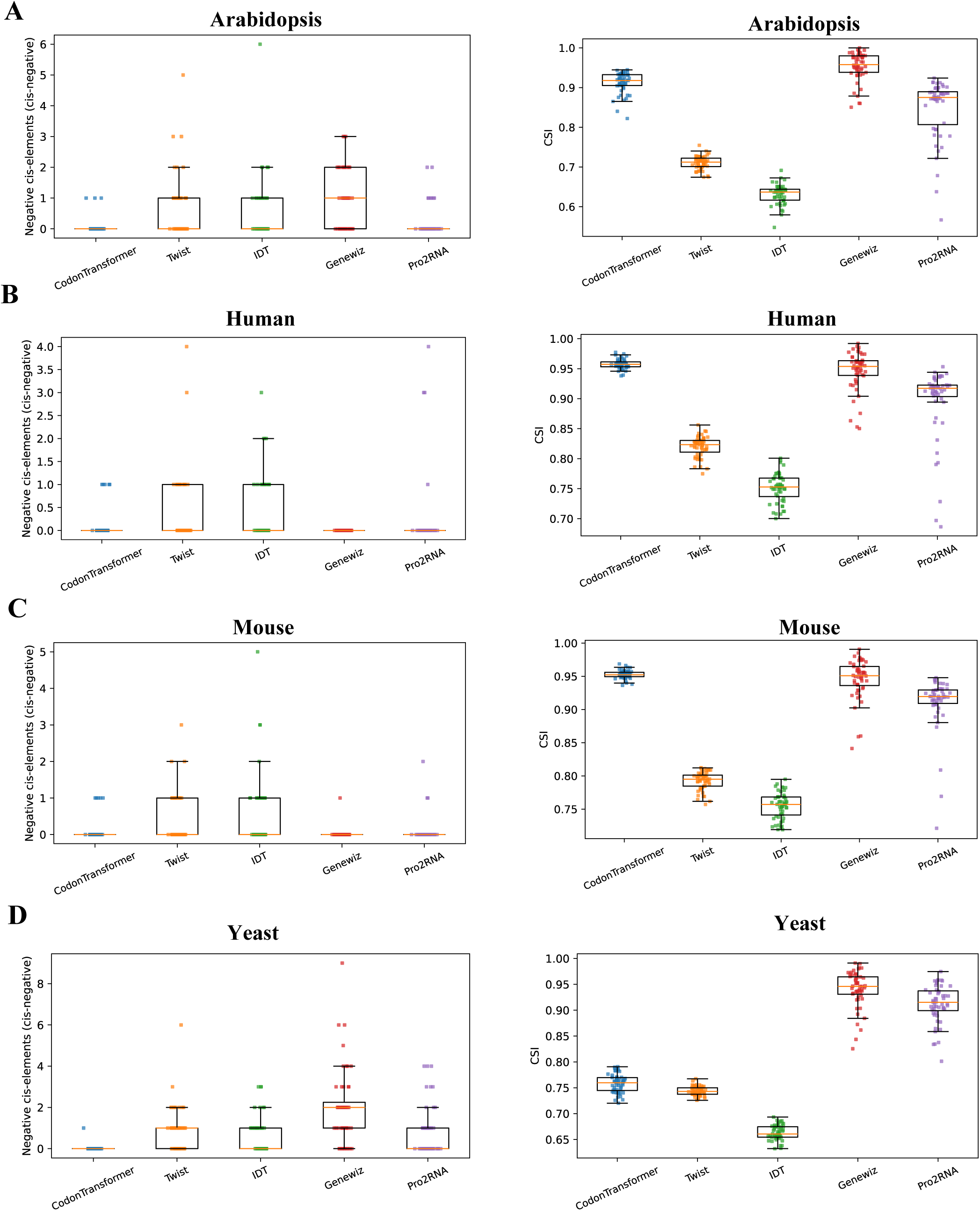
Pro2RNA-eukaryote generates coding sequences optimized for heterologous expression across eukaryotic hosts. Evaluation of heterologous coding sequence optimization in four eukaryotic model organisms: (A) Arabidopsis thaliana, (B) Human, (C) Mouse, and (D) Yeast. Left panels show the distribution of negative cis-regulatory elements (CIS-negative) within generated coding sequences, reflecting potential inhibitory features for expression. Right panels report the Codon Similarity Index (CSI), measuring the similarity of optimized sequences to native host coding patterns.

In addition, we benchmarked Pro2RNA against the method used in the GEMORNA study by evaluating mRNA coding sequences designed for the same four protein targets reported in the GEMORNA paper (Zhang et al. 2025a). We found that both GEMORNA and Pro2RNA generated coding sequences with almost no negative *cis*-regulatory elements (Table S3). However, Pro2RNA consistently produced sequences with lower Codon Similarity Index (CSI) compared to those generated by GEMORNA. In contrast, GEMORNA-generated coding sequences exhibited CSI values close to 1, indicating strong signatures of artificial codon sequence design. By comparison, Pro2RNA-generated sequences showed CSI distributions that were closer to those of native host genes, suggesting a higher degree of naturalness and better adaptation to the host translational landscape (Table S3).

Taken together, these findings demonstrate that Pro2RNA learns a context-dependent codon encoding strategy that prioritizes functional expressibility over maximal codon bias. By jointly minimizing negative *cis*-regulatory elements and avoiding excessive codon over-optimization, Pro2RNA provides a robust and generalizable framework for heterologous mRNA coding design across diverse biological hosts.

## Discussion

In this study, we introduced Pro2RNA, a multimodal reverse-translation language modelling for generating host-specific mRNA coding sequences from protein sequences and species taxonomy information. The core architectural advantage of Pro2RNA lies in the integration of SciBERT, ESM-2, and generative RNA language model, which jointly capture complementary biological information at different semantic levels. SciBERT encodes taxonomy and biological text semantics, ESM-2 provides rich protein sequence representations, and generative RNA language model enables autoregressive generation of mRNA sequences. Ablation studies demonstrate that SciBERT, ESM2 and mRNA-GPT combination consistently outperforms alternative variants, indicating that biological text semantics and protein sequence features provide complementary guidance for mRNA generation. Notably, training across broad taxonomic coverage (“border species”) further improves performance compared with single-species models and yields strong generalization on unseen datasets, suggesting that the model learns taxonomy-aware and natural-like mRNA coding patterns.

To leverage large-scale curated datasets, we trained Pro2RNA-bacteria on approximately 1 million protein-mRNA pairs from 241 bacterial species and Pro2RNA-eukaryote on 0.55 million pairs from 13 eukaryotic species. Across both domains, Pro2RNA consistently outperforms or comparable existing academic and commercial methods in multiple evaluation metrics, including codon similarity, CSI, and DTW distance. These results indicate that Pro2RNA effectively captures both local and global coding patterns characteristic of native host organisms.

In heterologous expression tasks, Pro2RNA learns organism-specific genetic codes and coding patterns. Rather than aggressively maximizing codon optimality, Pro2RNA generates mRNA coding sequences with intermediate CSI values and reduced numbers of negative *cis*-regulatory elements. This balanced behavior is biologically meaningful, as accumulating evidence suggests that optimal gene expression does not arise from maximal matching between codon usage and cellular tRNA supply (Chen et al. 2026). Instead, a slight mismatch between codon usage bias (CUB) and tRNA availability may be evolutionarily maintained to balance functional payoff with translational cost (Chen et al. 2026). Excessive codon optimization can increase mRNA secondary structure, disrupt beneficial ribosome pausing, impair co-translational protein folding, and potentially impose unnecessary translation burden (Yu et al. 2015; Liu 2020). Consistent with recent findings that intermediate CUB–tRNA mismatch can confer higher fitness than either minimized or excessive mismatch (Chen et al. 2026), Pro2RNA appears to implicitly capture this evolutionary trade-off. Together, these properties suggest that mRNA coding sequences generated by Pro2RNA are more aligned with natural selection principles and thus more suitable for practical heterologous expression in real biological systems.

Beyond predictive performance, Pro2RNA offers a flexible and resource-efficient design paradigm for mRNA coding sequence generation. Its modular architecture allows individual components: text, protein, or RNA language models, to be replaced or upgraded as improved models become available. Moreover, parameter-efficient fine-tuning enables rapid adaptation to new host organisms using user-provided protein-mRNA pairs and taxonomy information, without retraining the full model from scratch. This compositional strategy combines the strengths of specialized pretrained models while maintaining computational efficiency, making Pro2RNA a practical and extensible framework for rational mRNA coding sequence design.

In conclusion, compared with existing academic and commercial codon optimization tools, Pro2RNA consistently achieves superior or comparable results in both native sequence recovery and heterologous expression tasks, highlighting its potential as a general-purpose for rational mRNA coding sequence design.

## Material and Method

### Dataset collection and curation

Taxonomic annotations, mRNA coding sequences, and corresponding protein sequences were downloaded from NCBI using the NCBI Datasets command-line tools. To construct high-quality mRNA-protein pairs, we applied an in-house validation pipeline to ensure that each mRNA coding sequence exactly encodes its associated protein sequence, excluding incomplete, inconsistent, or mismatched entries. For training Pro2RNA-bacteria, we collected approximately 1 million mRNA-protein pairs from 241 bacterial species. For training Pro2RNA-eukaryote, we curated approximately 0.55 million mRNA-protein pairs from 13 representative eukaryotic species. All sequences were standardized to RNA format (with thymine replaced by uracil), and only protein-coding transcripts with unambiguous start and stop codons were retained for downstream modeling.

### Model architecture

We propose Pro2RNA, a modular and parameter-efficient framework for mRNA coding sequence generation conditioned on both protein sequence and host taxonomy information (Figure 1B). Pro2RNA consists of three pretrained language models integrated through feature fusion layers:

#### 1. Taxonomy encoder

Host taxonomic information is encoded using a pretrained scientific language model (SciBERT), which captures semantic representations of taxonomic descriptors. The taxonomy information is represented as a natural language prompt describing the hierarchical taxonomic classification (e.g., “The organism [name] belongs to the order [order], family [family], genus [genus], and species [species]”). This textual representation is tokenized and embedded using the SciBERT tokenizer, producing a single vector representation that captures the taxonomic context.

#### 2. Protein encoder

The target protein sequence is encoded using a pretrained protein language model (ESM-2), which provides rich residue-level representations learned from large-scale protein sequence corpora. The protein sequence is tokenized using the ESM-2 tokenizer, and the final layer representations are extracted to obtain contextualized residue-level embeddings.

#### 3. Generative RNA decoder

The mRNA coding sequence is generated using either a GPT-style autoregressive language model (mRNA-GPT or generRNA) or a simple MLP decoder, depending on the configuration. The decoder operates at the codon level, generating one codon token at a time in an autoregressive manner.

To efficiently adapt large pretrained models while minimizing trainable parameters, Low-Rank Adaptation (LoRA) is applied to all three components (Hu et al. 2022). The parameters of SciBERT, ESM-2, and the RNA decoder (mRNA-GPT or generRNA) are frozen, while only the LoRA adapters and the feature fusion layers are updated during training.

The outputs of the taxonomy encoder and protein encoder are fused through feature concatenation followed by multi-layer perceptron (MLP) projection. Specifically, the species embedding (a single vector) is expanded to match the sequence length of protein embeddings and concatenated along the feature dimension. The concatenated features are then projected through a two-layer MLP to produce unified representations that encode both protein sequence features and host-specific taxonomic context. This design allows the generative RNA model to jointly leverage protein sequence information and organism-specific codon usage patterns.

### Model input

#### 1. Taxonomy information

Host taxonomy is provided as a short textual description of the organism’s taxonomic classification (e.g., domain, phylum, order, and family). The taxonomy text is encoded using a pretrained SciBERT model, and the resulting pooled representation is used as a species-level embedding to condition mRNA coding sequence generation.

#### 2. Protein sequence

Protein sequences are represented using standard single-letter amino acid codes. The sequences are encoded using a pretrained ESM-2 protein language model, which produces contextualized residue-level representations for downstream mRNA generation.

#### 3. mRNA sequence

The target mRNA sequence is represented at the codon level using a minimal codon tokenizer with a vocabulary of 68 tokens: 64 standard codons (including the three stop codons UAA, UAG, and UGA) and four special tokens (<cls>, <pad>, <eos>, <unk>). Each codon is treated as a discrete token, and sequences are tokenized by splitting the mRNA sequence into non-overlapping 3-nucleotide codons. During training, the model learns to autoregressively predict codon tokens conditioned on both protein and taxonomy embeddings.

### Model training

#### Training objective

Pro2RNA is trained using an autoregressive next-token prediction objective on the mRNA coding sequence. Given a protein sequence and host taxonomy, the model learns to maximize the conditional likelihood:

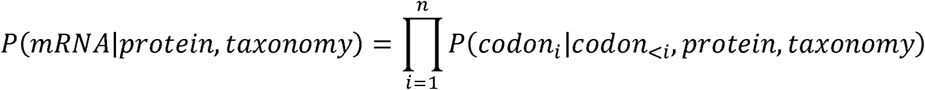

where *codon_i_* represents the i-th codon in the mRNA sequence. Cross-entropy loss is computed between the predicted logits and ground-truth codon token indices, with padding tokens ignored during loss computation. The loss is averaged over all non-padding positions in the sequence.

#### Training strategy

To achieve parameter-efficient training, all backbone parameters of SciBERT, ESM-2, and the RNA decoder (mRNA-GPT or generRNA) are frozen. Trainable parameters include: (1) LoRA adapters applied to the attention layers (query, key, value, and output projections) in SciBERT and ESM-2, and to the attention and feed-forward layers in the RNA decoder; (2) MLP projection layers that fuse protein and taxonomy embeddings; (3) output layers (language model head) for codon prediction. This design significantly reduces memory consumption and stabilizes training while maintaining the representational capacity of the pretrained models.

#### Optimization details

Optimization is performed using the AdamW optimizer with default hyperparameters. Training is conducted on eight NVIDIA H200 GPUs using data parallelism. The learning rate follows a cosine annealing schedule with warm-up steps. Gradient accumulation is employed to effectively increase the batch size while maintaining memory efficiency. Early stopping and checkpoint selection are based on validation loss, with the best model checkpoint saved based on the lowest validation loss. This parameter-efficient training strategy allows Pro2RNA to scale to millions of protein-mRNA pairs while maintaining stable convergence and requiring minimal computational resources compared to full fine-tuning approaches.

#### Inference

During inference, Pro2RNA generates mRNA coding sequences in an autoregressive manner. Host taxonomy prompts are first encoded by a pretrained SciBERT model, while the input protein sequence is encoded by a pretrained ESM-2 protein language model. Both encoders remain frozen, with lightweight LoRA adapters applied to enable task-specific adaptation. The resulting taxonomy and protein representations are concatenated and passed through a trainable MLP projector, which maps the fused representation into the latent space of the RNA decoder. This projected conditioning vector is then provided to the mRNA-GPT decoder, which is also frozen except for LoRA modules. Conditioned on the fused taxonomy-protein representation and previously generated tokens, the RNA decoder generates codons sequentially in an autoregressive fashion. Decoding is performed using greedy decoding or beam search, depending on the application scenario. The generation process terminates when a stop codon is produced or when the maximum sequence length is reached. This design enables species-aware, protein-conditioned mRNA coding sequence generation, making Pro2RNA suitable for applications such as heterologous expression, codon optimization, and host-specific mRNA design.

#### Calculation of negative cis-regulatory elements

The number of negative cis-regulatory elements is calculated using the Genescript codon analysis tool (https://www.genscript.com/tools/rare-codon-analysis), which was previously adopted in the CodonTransformer study.

#### Codon similarity score and naturalness score determination

To quantify positional agreement between a generated coding sequence and the native CDS encoding the same protein, we compute the codon similarity score and naturalness score as the fraction of codons that are identical at the same position. The codon similarity score is calculated for each individual mRNA coding sequence, whereas the naturalness score is computed at the dataset level, aggregating results across all mRNA coding sequences in the test set.

Given two codon sequences *A=(a_1_,a_2_,…,a_L_)*and *B=(b_1_,b_2_,…,b_L_)* with equal length *L*, the codon similarity score is defined as:

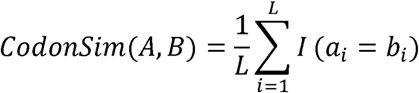

where *I(⋅)* is the indicator function (i.e., *I(a_i_=b_i_)*=1 if *a_i_=b_i_* and 0 otherwise).

#### Codon similarity index (CSI)

We evaluate codon-usage adaptation using the Codon Similarity Index (CSI) (Sabath et al. 2012), which quantifies how well a gene’s codon usage matches that of the host organism by computing the geometric mean of relative codon adaptiveness values across all coding positions. In this study, host codon usage frequencies from the Kazusa Codon Usage Database (Nakamura et al. 2000) are used as the reference.

For each amino acid, let *x_ij_* denote the observed usage frequency of codon *j* encoding amino acid *i* in the host genome. The relative adaptiveness of codon for *j* amino acid *i* is defined as

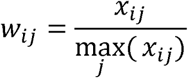

Given a sequence of length *L* codons (excluding stop codons), and denoting the relative adaptiveness of the codon at position *k* as *w_k_*, the CSI of this sequence is computed as the geometric mean:

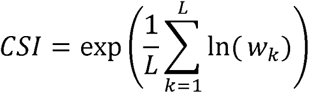

#### %MinMax profiles and DTW distance

Local codon-usage smoothness is assessed via %MinMax profiles (Clarke and Clark 2008), computed using a sliding window of 18 codons. For each window *i,* let *f*_c_ be the host usage frequency of codon *c* and *f̅_c_*, *f*_min,*c*_ and *f*_max,*c*_ denote the mean, minimum, and maximum frequencies among all synonymous codons for the same amino acid. In this section, codon usage frequencies are also from the Kazusa Codon Usage Database (Nakamura et al. 2000).

The %MinMax value for window *i* is then defined as

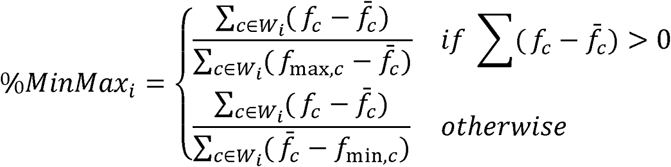

To compare generated and native sequences, we compute the Dynamic Time Warping (DTW) distance (Müller 2007; Fallahpour et al. 2025) between their %MinMax

Given two profiles *X(X_1,…,_X_m_*) and *Y=(Y_1,..,_Y*_n_), the DTW distance is defined as

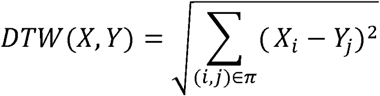

where *π* denotes the optimal alignment path that minimizes the cumulative squared deviation.

## Code availability

The codes used for Pro2RNA training and inference can be found at https://github.com/ZHymLumine/Pro2RNA/

## Acknowledgements

This work was supported by JST CREST JPMJCR23N1. The computations were partially performed on the NIG supercomputer at ROIS National Institute of Genetics, the ABCI supercomputer at AIST, HOKUSHIN supercomputer at Department of Data Science, Kitasato University and the SQUID supercomputer at Cybermedia Center, Osaka University through the HPCI System Research Project (hp230057, hp240075).

---

**Figure S1.**
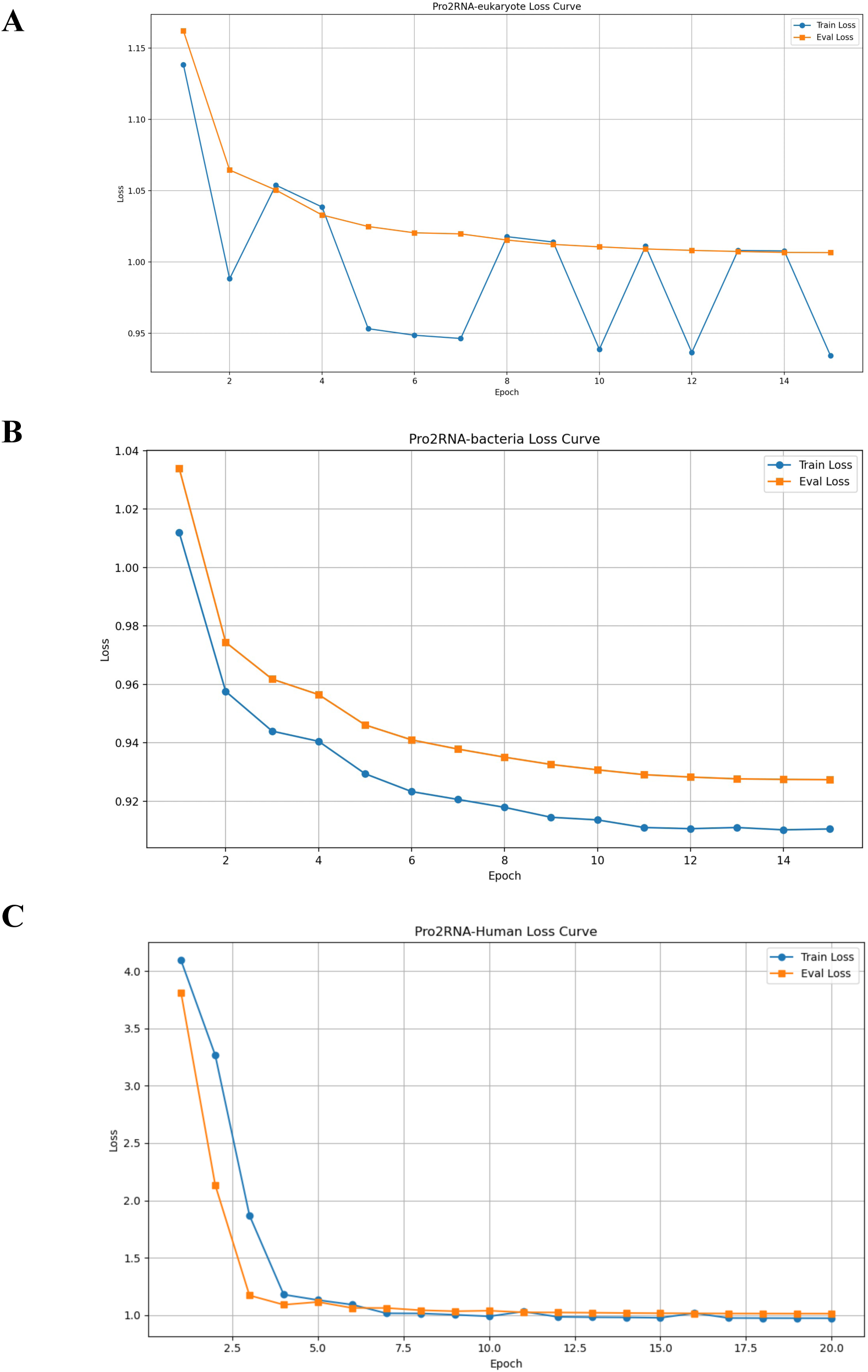
Loss curve of training Pro2RNA-eukaryote (A), Pro2RNA-bacteria (B) and Pro2RNA-human (C).

**Table S1.** Eukaryotic species used for training the Pro2RNA-eukaryote model. List of representative eukaryotic species whose mRNA coding sequences were used to train the Pro2RNA-eukaryote model. For each species, the corresponding RefSeq genome assembly ID and scientific name are provided. The selected organisms span diverse eukaryotic lineages, including mammals, insects, plants, fungi, and protists, ensuring broad taxonomic coverage for learning species-specific and lineage-dependent codon usage patterns.

| RefSeq ID | Scientific Name |
| --- | --- |
| GCF_000001635.27 | *Mus musculus* (house mouse) |
| GCF_000001405.40 | *Homo sapiens* (human) |
| GCF_036323735.1 | *Rattus norvegicus* (Norway rat) |
| GCF_000001215.4 | *Drosophila melanogaster* (fruit fly) |
| GCF_000002035.6 | *Danio rerio* (zebrafish) |
| GCF_000146045.2 | *Saccharomyces cerevisiae* S288C |
| GCF_000002945.2 | *Schizosaccharomyces pombe* (fission yeast) |
| GCF_000002985.6 | *Caenorhabditis elegans* |
| GCF_000001735.4 | *Arabidopsis thaliana* (thale cress) |
| GCF_034140825.1 | *Oryza sativa* Japonica Group (Japanese rice) |
| GCF_000027005.1 | *Komagataella phaffii* GS115 |
| GCF_000004695.1 | *Dictyostelium discoideum* |
| GCF_000002765.6 | *Plasmodium falciparum* 3D7 |

**Table S2.** Bacterial species used for training the Pro2RNA-bacteria model. Summary of bacterial species included in the training dataset of the Pro2RNA-bacteria model, grouped by taxonomic family. For each family, the number of representative organisms is listed. The selected families cover diverse bacterial clades with distinct genomic and codon usage characteristics, ensuring balanced representation and enabling the model to learn family- and species-specific coding patterns.

| Family | Number of Organisms |
| --- | --- |
| Bacillaceae | 40 |
| Bacteroidaceae | 40 |
| Enterobacteriaceae | 60 |
| Mycobacteriaceae | 40 |
| Pseudomonadaceae | 41 |
| Streptococcaceae | 40 |

**Table S3.** Evaluation of heterologous coding sequence optimization in human compared with GEMORNA. Comparison of coding sequences optimized by Pro2RNA and GEMORNA for multiple heterologous proteins expressed in human, including Firefly luciferase, SARS-CoV-2 spike protein, erythropoietin (EPO), NanoLuc, and chimeric antigen receptor (CAR) constructs. For each protein, multiple independently generated sequences are reported. Performance is assessed using the Codon Similarity Index (CSI) and the presence of negative cis-regulatory elements (cis-negative).

| Protein | Method | CSI | cis-negative |
| --- | --- | --- | --- |
| Firefly luciferase | GMR-FL1 | 0.98773967 | 0 |
| Firefly luciferase | GMR-FL2 | 0.97999307 | 0 |
| Firefly luciferase | GMR-FL3 | 0.94818749 | 0 |
| Firefly luciferase | GMR-FL4 | 0.92789634 | 0 |
| Firefly luciferase | Pro2RNA-FL1 | 0.80313026 | 0 |
| Firefly luciferase | Pro2RNA-FL2 | 0.79972448 | 0 |
| Firefly luciferase | Pro2RNA-FL3 | 0.79086332 | 0 |
| Firefly luciferase | Pro2RNA-FL4 | 0.79197337 | 0 |
| Covid-19 spike | GMR-CV1 | 0.96138753 | 0 |
| Covid-19 spike | GMR-CV2 | 0.98744619 | 0 |
| Covid-19 spike | GMR-CV3 | 0.91723779 | 0 |
| Covid-19 spike | Pro2RNA-CV1 | 0.76904283 | 1 |
| Covid-19 spike | Pro2RNA-CV2 | 0.7716272 | 0 |
| Covid-19 spike | Pro2RNA-CV3 | 0.77022572 | 0 |
| EPO | GMR-EPO1 | 0.89136957 | 0 |
| EPO | GMR-EPO2 | 0.9335418 | 0 |
| EPO | GMR-EPO3 | 0.9003293 | 0 |
| EPO | GMR-EPO4 | 0.91560663 | 0 |
| EPO | GMR-EPO5 | 0.90012195 | 0 |
| EPO | Pro2RNA-EPO1 | 0.85316816 | 0 |
| EPO | Pro2RNA-EPO2 | 0.80121476 | 0 |
| EPO | Pro2RNA-EPO3 | 0.80961562 | 0 |
| EPO | Pro2RNA-EPO4 | 0.80144251 | 0 |
| EPO | Pro2RNA-EPO5 | 0.87003975 | 0 |
| NanoLuc | GMR-NL1 | 0.97925621 | 0 |
| NanoLuc | GMR-NL2 | 0.98580451 | 0 |
| NanoLuc | GMR-NL3 | 0.9291135 | 0 |
| NanoLuc | GMR-NL4 | 0.94883676 | 0 |
| NanoLuc | GMR-NL5 | 0.93915275 | 0 |
| NanoLuc | Pro2RNA-NL1 | 0.80479587 | 0 |
| NanoLuc | Pro2RNA-NL2 | 0.75325533 | 0 |
| NanoLuc | Pro2RNA-NL3 | 0.75858341 | 0 |
| NanoLuc | Pro2RNA-NL4 | 0.78121929 | 0 |
| NanoLuc | Pro2RNA-NL5 | 0.79014909 | 0 |
| CD19 CAR | GMR-CD19-1 | 0.96797707 | 0 |
| CD20 CAR | GMR-CD19-2 | 0.97122968 | 0 |
| CD21 CAR | Pro2RNA--CD19-1 | 0.86352859 | 0 |
| CD22 CAR | Pro2RNA--CD19-2 | 0.87015268 | 0 |

